# Breast cancer mutations HER2^V777L^ and PIK3CA^H1047R^ activate the p21-CDK4/6 –Cyclin D1 axis driving tumorigenesis and drug resistance

**DOI:** 10.1101/2022.11.09.515796

**Authors:** Xiaoqing Cheng, Yirui Sun, Maureen Highkin, Nagalaxmi Vemalapally, Xiaohua Jin, Brandon Zhou, Julie L. Prior, Ashley R. Tipton, Shunqiang Li, Anton Iliuk, Samuel Achilefu, Ian S. Hagemann, John R. Edwards, Ron Bose

## Abstract

In metastatic breast cancer, HER2 activating mutations frequently co-occur with mutations in the *PIK3CA*, *TP53*, or E-cadherin genes. Of these co-occurring mutations, *HER2* and *PIK3CA* mutations are the most prevalent gene pair, with approximately 40% of *HER2* mutated breast cancers also having activating mutations in *PIK3CA*. To study the effects of co-occurring *HER2* and *PIK3CA* mutations, we bred genetically engineered mice with the *HER2^V777L^*; *PIK3CA^H1047R^*transgenes (HP mice) and studied the resulting breast cancers both *in vivo* as well as *ex vivo* using cancer organoids. HP breast cancers show accelerated tumor formation *in vivo* and increased invasion and migration in *in vitro* assays. HP breast cancers have resistance to the pan-HER tyrosine kinase inhibitor, neratinib, but are effectively treated by neratinib plus trastuzumab deruxtecan. Proteomic and RNA-Seq analysis of HP breast cancers showed increased gene expression of Cyclin D1 and p21WAF1/Cip1 and changes in cell cycle markers. Combining neratinib with CDK4/6 inhibitors was another effective strategy for HP breast cancers with neratinib plus palbociclib showing a statistically significant reduction in mouse HP tumors as compared to either drug alone. We validated both the neratinib plus trastuzumab deruxtecan and neratinib plus palbociclib combinations using a human breast cancer patient-derived xenograft that has very similar HER2 and *PIK3CA* mutations. Both of these drug combinations are being tested in phase 1 clinical trials and this study provides valuable preclinical evidence for them.

## Introduction

The HER2 receptor tyrosine kinase is a major therapeutic drug target in breast cancer. Trastuzumab and other HER2 targeted drugs have dramatically improved outcomes for patients with HER2 gene amplified breast cancers (1). One of the major findings of The Cancer Genome Atlas (TCGA) Breast Cancer project was the identification of HER2 activating mutations in breast cancers without HER2 gene amplification (2,3). These HER2 mutations are clustered missense mutations either in the tyrosine kinase domain or at codons 309-310 of the extracellular domain (3). The HER2^V777L^ mutation in the kinase domain is a particularly strong mutation, showing 20-fold increase in HER2 *in vitro* kinase activity and strong effects in cell-based assays (3). Therefore, we selected the HER2^V777L^ mutation for creation of a novel transgenic mouse (4).

HER2 activating mutations are very sensitive to the oral pan-HER tyrosine kinase inhibitor, neratinib, in preclinical models. Clinical trials have tested neratinib monotherapy or the combination of neratinib with the hormonal therapy drug, fulvestrant, to treat metastatic breast cancer (MBC) patients whose cancer carry HER2 mutations. These clinical trials have found clinical benefit in 28 to 46% of HER2 mutated MBC patients but median progression-free survival was only 3.6 to 5.4 months (5-9). Co-occuring mutations in *TP53,* HER3, and *PIK3CA* genes have been implicated as factors in patients who do not benefit from neratinib (5,10). Of these, *PIK3CA* mutations are the most common, with approximately 40% of HER2 mutated metastatic breast cancers also containing activating mutations in *PIK3CA (6)*. *PIK3CA* mutations have been linked to drug resistance in HER2 gene amplified breast cancer (11,12) prompting us to study the combination of HER2 and *PIK3CA* mutations in a preclinical model system. The availability of *PIK3CA*^H1047R^ transgenic mice provides a way to study this pair of activating mutations in a genetically engineered mouse model (13). Mice with *PIK3CA*^H1047R^ and *TP53* mutations develop adenosquamous carcinoma, adenomyoepitheliomas, and spindle/EMT tumors in the mammary glands whereas mice with *PIK3CA*^H1047R^ alone predominantly developed fibroadenomas with a few adenocarcinomas or spindle cell neoplasms (14,15). Mice with *PIK3CA*^H1047R^ plus E-cadherin deletion develop invasive lobular carcinoma of breast (16). *PIK3CA^H1047R^* mutation enhanced tumor formation in wild-type, human HER2 transgenic mice and generated drug resistance to the HER2 targeted drugs, trastuzumab, pertuzumab, and lapatinib (11).

Organoid culture is a 3D culture cell technique that mimics the *in vivo* cell niche and increases the success rate for growing primary cell cultures. Organoids models are increasingly used in translational cancer research, as they accelerate functional studies and drug testing on cancer samples (17). Human patient-derived xenografts (PDXs) and matched PDX-derived organoids (PDxO) have been validated for precision oncology (18). Breast cancer organoids can be grown long term and recapitulate the complex genetic and phenotypic heterogeneity of breast cancers (19). In this study, we isolated organoids from genetically-engineered mouse breast cancers and used them for preclinical therapeutic and mechanistic experiments.

This paper reports the generation of HER2^V777L^; *PIK3CA^H1047R^* transgenic mice (HP mice) and HP mutated breast cancer organoids to study tumor formation, drug sensitivity, gene expression changes and cell cycle alterations in breast cancers that have mutations in both of these genes. We found that the HP mutation combination activates the p21-CDK4/6-CyclinD1 axis to drive tumorigenesis and drug resistance.

## Results

### Tumor formation in mice with combined HER2^V777L^ and *PIK3CA^H1047R^* expression

We previously reported the development of a novel transgenic mouse that conditionally expresses the human HER2 V777L cDNA (which we will abbreviate as “H”), which is inserted into the Rosa26 locus using TALEN-based genome editing (4). Our HER2 mutant transgene has a lox-STOP-lox cassette located immediately 5’ to the transgene, and the transgene will only express after Cre recombinase-mediated removal of the lox-STOP-lox cassette (Fig. 1A). This transgenic mouse, Rosa26-LSL-human HER2 ^V777L^ (H), was bred with Rosa26-LSL-*PIK3CA ^H1047R^* mice (which we will abbreviated as P) to create the HP mouse as shown in Fig. 1B (14). *PIK3CA^H1047R^* is a gain-of-function allele and activating mutation that is commonly found in human breast cancers. Both the *PIK3CA^H1047R^* transgene and our HER2^V777L^ mutant transgene were inserted into the Rosa26 locus of the mouse genome, and therefore mice with both transgenes (HP mice) have one copy of each transgene (Fig. 1B).

**Fig. 1:**
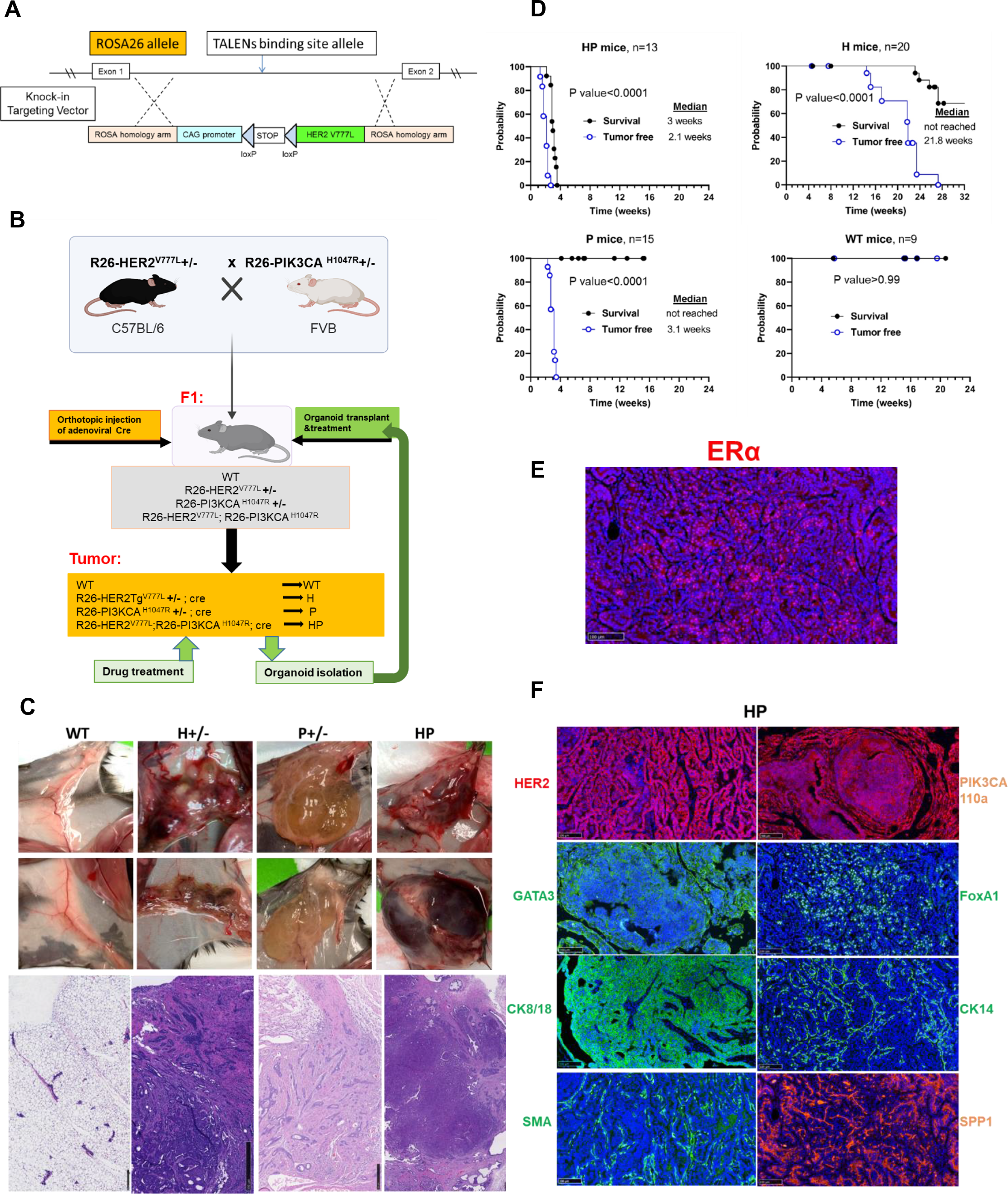
Tumor formation in HER2^V777L^ and *PIK3CA^H1047R^* transgenic mice. (A) Schematic for design of HER2^V777L^ Tg mouse. (B) schematic view of wide type (WT), HER2^V777L^ (H), *PIK3CA*^H1047R^ (P), HER2^V777L^; *PIK3CA*^H1047R^ (HP) transgenic mouse model. (C) Top, representative brightfield image showing the primary tumor in the mammary gland induced by Cre recombination using adenovirus injection. Bottom, representative hematoxylin and eosin (H&E) staining of the tumor-bearing mammary gland of wide type, H, P, HP animals after Cre adenovirus injection. (D) Kaplan-Meier analysis shows overall survival (black line) and tumor-free (blue line) of WT, H, P, and HP group mice respectively. (E) IF staining of ERα images of mammary gland tumor tissue sections from HP mice. Scale bars are 100 µm. (F) immunofluorescence (IF) staining of HER2, PIK3CA, SPP1, GATA3, SMA, FOXA1, CK14, CK8/18 images of mammary gland tumor tissue sections from HP mice. Scale bars are 100 µm.

Mammary tumors were generated in these mice by orthotopic injection of adenovirus expressing Cre (Ad-Cre) into 10 weeks old female mice (Fig. 1C). H mice slowly developed areas of *in situ* carcinomas and invasive adenocarcinomas over a 4–6-month period after Ad-Cre injection (Fig. 1C, D). P mice developed tumors with scirrhous growth patterns, as well as cystic areas resembling lymphangiomas, as previously described (14,16). HP mice rapidly developed invasive mammary adenocarcinoma (Fig. 1C). Remarkably, HP mice rapidly developed breast cancers at a median time of 2.1 weeks post Ad-Cre injection and had to be euthanized at 3 weeks due to large tumor size (Fig. 1D). In contrast, P mice formed cystic tumors at a median time of 3.1 weeks, but the tumors did not kill the mice over a 16 weeks’ time frame. H mice developed palpable tumors at a median time of 21.8 weeks and 65% of H mice are alive at 28 weeks (Fig. 1D).

Multiple immunostains were performed on the adenocarcinomas from HP mice to characterize the mammary epithelial lineage and expression patterns of HP tumors. The HP tumors are ER positive, HER2 positive, and PIK3CA positive (Fig 1E, F). Expression of the key ER regulators GATA3 and FoxA1 demonstrates HP tumors derive from mammary epithelial cells. Positive cytokeratin 8/18 and negative cytokeratin 14 and smooth muscle actin (SMA) indicate that these are luminal type cells, rather than basal cells (Fig 1F). Osteopontin (OPN) and *SPP1* mRNA expression are related to recurrence in ER positive breast cancer (20). Positive OPN expression indicate HP tumors may have a high risk of recurrence and improved therapies for them are needed.

### *In vitro* invasion with organoids derived from HER2^V777L^ expression and *PIK3CA*^H1047R^ tumors

We isolated breast organoids from WT, H, P, HP mice (Fig. 2A) in order to obtain an enriched epithelial cell population. To assess the cellular migration and invasion signature of HER2^V777L^ and *PIK3CA^H1047R^* breast epithelium *in vitro*, we performed branching morphogenesis and invasion assays with the breast organoid cultures. For the branching morphogenesis assay, 2.5nM FGF2 was added to both P and HP organoids, resulting in an increased number of buddings in only the HP organoids (Fig. 2B). Additionally, we assessed the cell invasion assay by adding FGF2 to organoids embedded in collagen I gel, and we observed an increased number of elongated buddings in both P and HP organoids (Fig. 2B). We next assessed the mobility and invasiveness of the organoids through transwell cell migration and invasion assays. Remarkably, the HP organoids showed increased migration and invasion into the surrounding extracellular matrix compared to the P organoids (Fig. 2C and 2E). These data suggest that the combination of HP mutations promote cancer cell invasion and migration. To confirm the enhanced cellular migration and invasion in HP organoids, we also performed a wound-healing assay to monitor the healing speed of the epithelial monolayer under the microscope every 4 hours (Fig. 2D). Organoids derived from HP mice tumors were observed to heal faster post scratching (Fig. 2D) as compared to organoids derived from the WT, H, and P mice. These data shows that HP organoids show increased cell migration and invasion, which are properties of metastatic breast cancer.

**Fig. 2:**
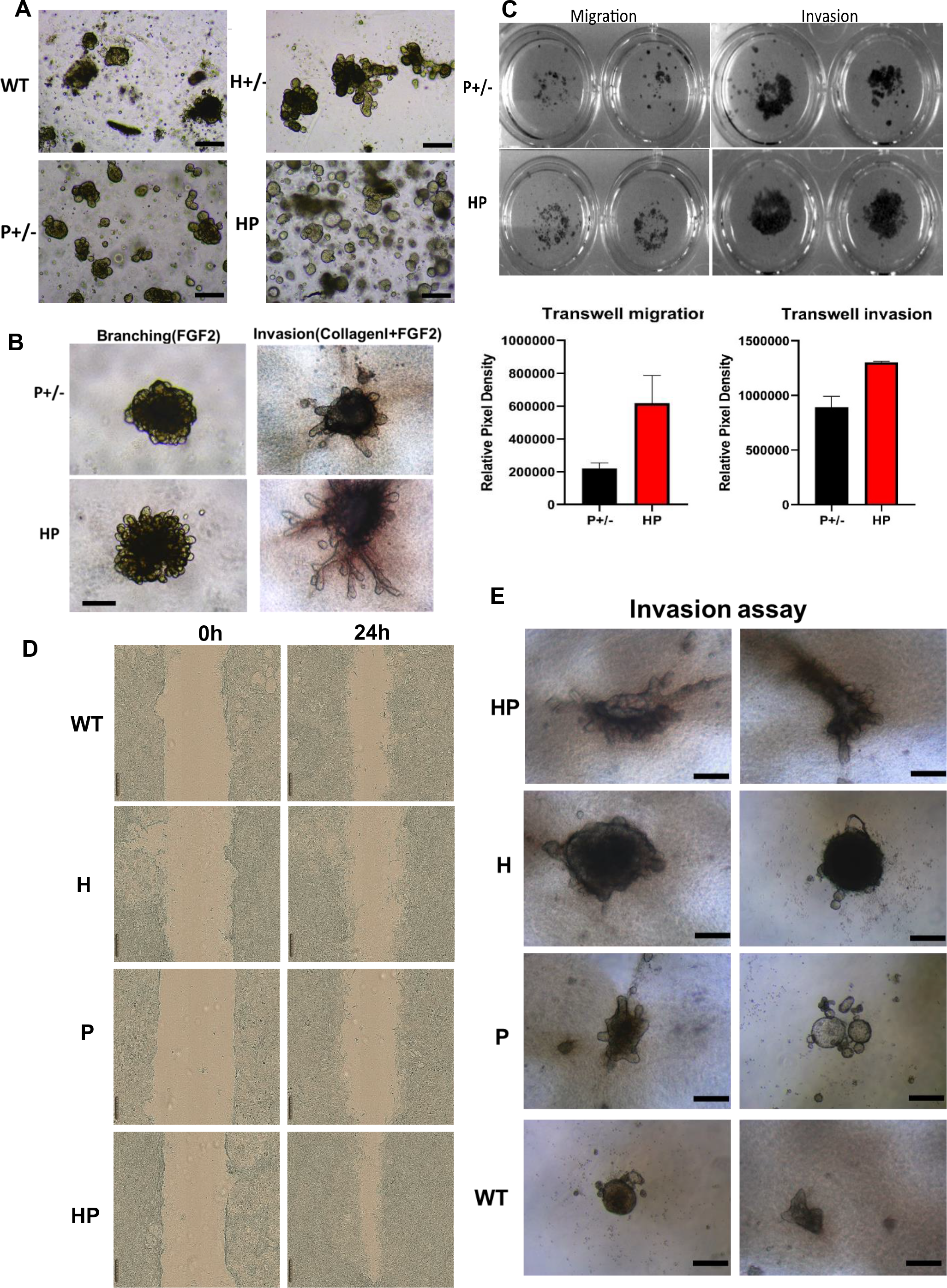
*In vitro* invasion with organoids derived from HER2^V777L^ and *PIK3CA^H1047R^* tumors. (A) representative image of organoid derived from mammary gland tissues from the WT, H, P, and HP mice. (B) Branching and invasion assay of organoid derived from P and HP mice tumor. Representative images are shown. Scale bars are 500 µm. (C) Transwell migration and invasion assay of organoid derived from P and HP mice tumor. Representative images are shown. Quantification of relative pixel density was calculated by ImageJ. (D) Wound-healing assay to monitor the healing speed of the epithelial monolayer under the microscope every 4 hours using incucyte S3. Scale bar = 100 µm. Original magnification × 10. (E) Invasion assay of organoid derived from H, P and HP mice tumor and WT mice. Representative images are shown. Scale bars are200 µm.

### Lung metastases in HER2^V777L^ transgenic mice and HP organoid transplant mice

We hypothesized that the HER2^V777L^ mutation plays a vital role in the metastatic breast cancer. To test this, H, P and HP mice were examined for metastasis to the lung or liver. Metastasis were not seen in HP mice likely because of the rapid growth of the primary tumor leading to early euthanasia. In P mice, no metastases were seen at 4 months after adenovirus cre injection into the mammary glands. Lung metastasis was observed in 85% of transgenic H mice (Fig. 3A), matching the invasive phenotype observed *in vitro* (Fig. 2). The histology, HER2 and ER staining of the lung metastasis matched the breast primary tumor (Fig. 3B). In order to improve visualization of both primary tumor and metastases, we conjugated trastuzumab with the near infrared (NIR) fluorophore, LS288 (21). H mice injected with this NIR-trastuzumab imaging agent show tumor-specific uptake at 24 hours in the breast and lung, but not in the liver or spleen (Fig. 3C). The imaging data suggested that the lung is a metastatic site for the HER2^V777L^ expressing breast tumor cells in our mouse models.

**Fig. 3:**
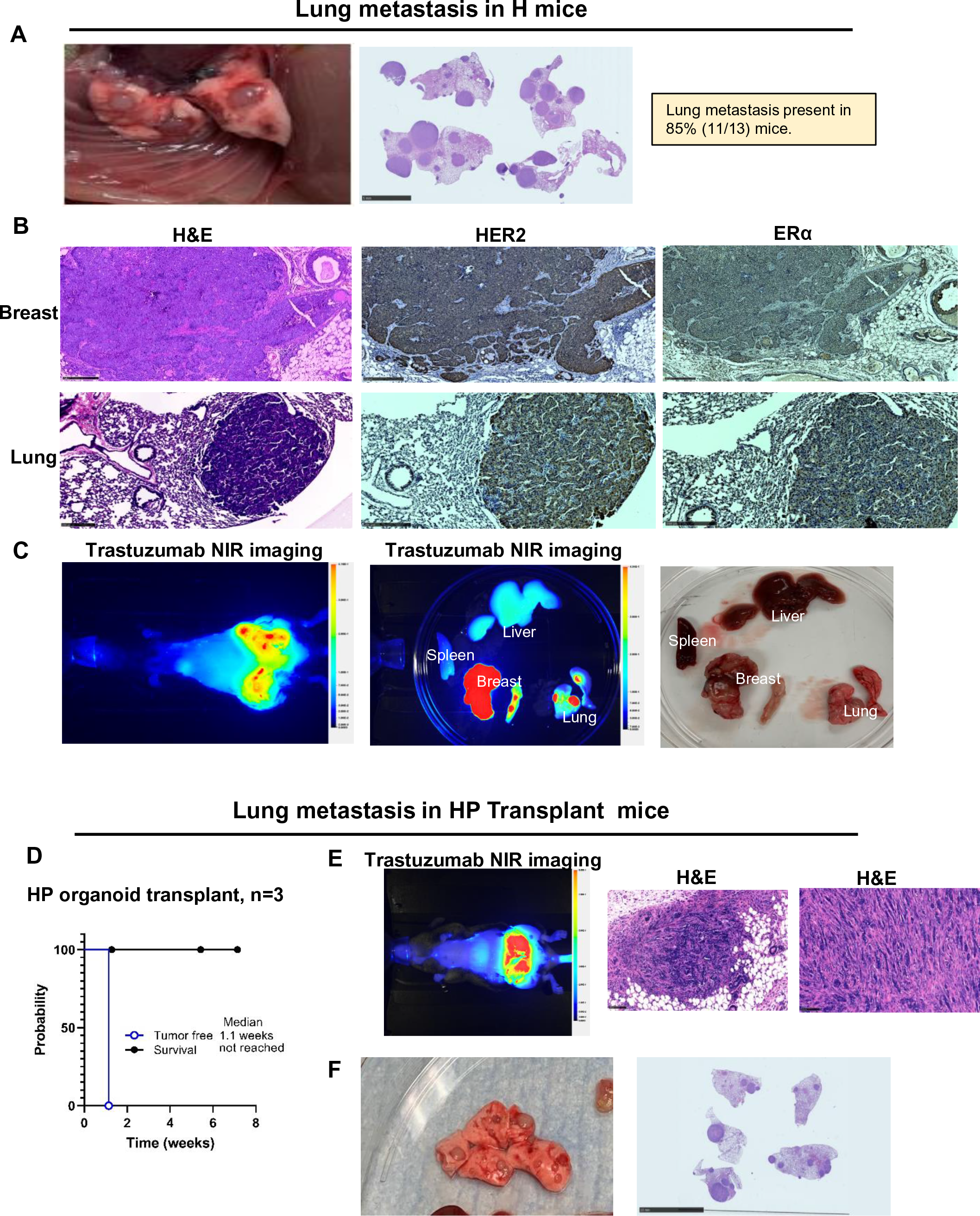
Lung metastases in HER2^V777L^ transgenic mice and HP organoid transplant mice. (A) First, Representative brightfield image of metastasis lung isolated from H mice. Middle, H&E staining on the metastasis lung isolated from H mice. Last, Quantification of the incidence of lung metastasis in dissected H mice. (B) H&E staining and IHC image on the mammary gland and lung tissue slides. Scale bars are 250 µm. (C)Top, Pearl imaging of NIR-trastuzumab at 24 hours post-injection on the whole body of H mice. Middle, Pearl imaging of NIR-trastuzumab at 24 hours post-injection on the mammary gland, spleen, lung, and liver tissues isolated from H mice. Last, Representative brightfield image of mammary gland, spleen, lung, and liver tissues isolated from H mice. (D) Kaplan-Meier analysis shows overall survival (black line) and tumor-free (blue line) of transplanted mice respectively. (E)Top, Pearl imaging of NIR-trastuzumab at 24 hours post-injection on the whole body of HP mice derived organoid transplanted mice. Middle and Last, Representative image of H&E staining on the breast tumor isolated from HP mice organoid transplanted mice. (F) Top, Representative brightfield image of metastasis lung isolated from HP mice organoid transplanted mice. Middle, H&E staining on the metastasis lung isolated from HP mice organoid transplanted mice.

To determine if the HP tumor-derived organoids recapitulate the metastatic process, organoids from the HP mice were orthotopically transplanted bilaterally into the 4^th^ mammary glands of syngeneic, recipient mice. Breast tumors were palpable one week after organoid transplantation (Fig. 3D), and these recipient mice survived longer than the HP mice injected with Ad-Cre (compare Fig. 3D and 1D). Hematoxylin and eosin staining of the transplanted tumor showed the same histological morphology as HP mouse tumors (Fig. 3E, F). As expected, the prolonged survival time provided time for the development of lung metastasis in the HP organoid transplant mice, suggesting that organoid transplants are a potential model to study the mechanisms and therapy of breast cancer metastasis.

### Drug testing *ex vivo* using tumor-derived organoids and *in vivo* on mice

We tested the impact of the P transgene on the neratinib drug sensitivity of HER2 mutated breast cancer organoids. *Ex vivo* cultures of HP tumor organoids were very resistant to neratinib with IC50 values >2 uM (Fig. 4A). In contrast, H tumor organoids and P tumor organoids were neratinib sensitive with IC50 values of 0.01 to 0.02 uM (Fig. 4A). These murine breast cancer organoids are grown in media supplemented with EGF. Since neratinib is a dual EGFR/HER2 inhibitor, P tumor organoids are still dependent on EGF in the media and are killed by neratinib. The sensitivity of these organoids to the PIK3CA inhibitor, alpelisib, was also tested. HP and P tumor organoids were equally sensitive to alpelisib (IC50 0.02 to 0.05 uM), whereas H tumor organoids were less sensitive with an IC50 of 0.5 uM (Fig. 4A). Next, we conducted *in vivo* drug treatments on our transgenic mice. Neratinib, trastuzumab, alpelisib and the antibody-drug conjugate, trastuzumab deruxtecan (T-DXd), were tested on HP mice. *In vivo*, neratinib or trastuzumab alone did not delay tumor formation or change the survival of HP mice (Fig. 4B). Alpelisib had a modest effect on tumor formation in HP mice (Fig. 4C). Dramatically, T-DXd abrogated tumor formation over a 5-week period, but the dose of 30mg/kg IV weekly times 3 doses was too toxic to the mice and mouse deaths were seen starting at week 5 (Fig. 4C).

**Fig. 4:**
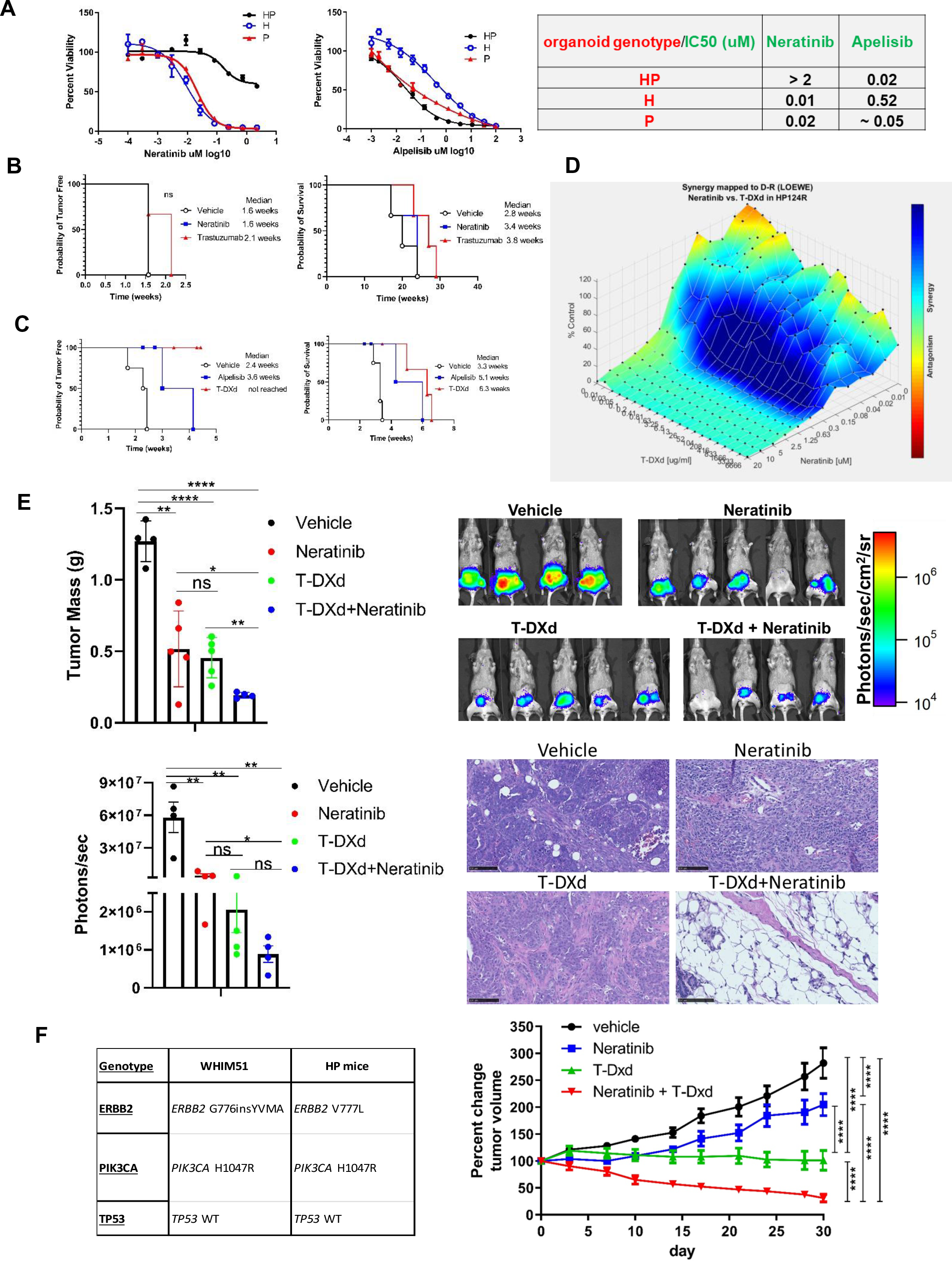
*Ex vivo* drug treatment using tumor-derived organoids and *in vivo* drug treatment on mice. (A) Cell viability for tumor organoids treated with neratinib or alpelisib for 6 days. (B) Kaplan-Meier analysis showing probability of tumor free and overall survival of HP mice treated with neratinib (40mg/kg daily by oral gavage) or trastuzumab (30mg/kg IV weekly x 3). Treatment was started 4 days post adenovirus-cre injection and n=3 per group. (C) Similar to (B). N=4 per group. Alpelisib dose was 25mg/kg daily by oral gavage and T-DXd was 30mg/kg IV weekly x 3 by tail vein injection. (D) Testing combined effect of T-DXd and neratinib. Drug synergy as per the Lowe model is indicated by blue colors on the 3D surface, whereas green colors indicate additivity. (E) HP breast tumor organoid cells (1 × 10^6^) were orthotopically injected into 8–10-week-old female FVB x C57BL/6J F1 (FVBB6F1) mice (n = 5–6). On day 4, starting the treatment with saline water, neratinib only, DS8201, and neratinib plus DS8201 respectively for 24 days. Neratinib was given daily at 40mg/kg by oral gavage. T-DXd was given weekly at 30mg/kg by tail vein injection for 2 doses. Top left, Tumors were dissected and weighed on day 28 post-injection upon collection. Top right, Representative BLI images of treated mice on day24 post-injection. Bottom left, BLI intensity measurements on day24 post injection. Bottom right, hematoxylin and eosin (H&E) staining of the tumor slides from transplant mice within treatment. Scale bars are 250 µm. (F) Left, comparison of the genotypes of human PDX WHIM51 and HP mice. Right, Tumor volume growth rate of treated mice in 4 arms as shown in the Figure. N=5 per arm. Data are plotted as means ± SEM. ∗P < .05, ∗∗P < .01, ∗∗∗P < .001, and ∗∗∗∗P < .0001 as calculated by the Mann-Whitney U test.

The combination of neratinib and T-DXd is being tested in two phase 1 clinical trials (ClinicalTrials.gov Identifiers NCT05372614 and NCT05274048) and therefore, evaluated this drug combination in HP mice. *Ex vivo,* we found strong synergy between neratinib and T-DXd (Fig. 4D). Neratinib has been shown to increase the internalization of trastuzumab containing ADC’s (22) and this is the likely mechanism for this drug synergy. Next, HP tumor organoids were luciferase labeled and transplanted into the mammary fat pads of syngenic mice. Luciferase imaging can be performed serial on mice without impacting tumor growth and is an accurate readout of tumor size (23). T-DXd was limited to two doses of 30mg/kg IV to avoid the mouse deaths previously seen at 5-6 weeks (Fig. 4C). The combination of neratinib plus T-DXd showed the greatest tumor shrinkage as measured by both tumor mass and luciferase signaling (Fig. 4E). Tumor shrinkage with this combination was statistically significant compared to vehicle and either drug on its own based on tumor mass measurements. A little more variation in the luciferase signal was seen in T-DXd treated mice and therefore, by luciferase signal measurement, statistical significance was seen between the neratinib + T-DXd combination and both vehicle and neratinib monotherapy, but not with T-DXd monotherapy. However, the histology of drug treated tumors shows a dramatic effect of the neratinib + T-DXd combination with clearance of tumor cells from the fat pad. This degree of tumor cell kill is not achieved with either T-DXd or neratinib monotherapy (Fig. 4E). Mice tolerated this combination of neratinib plus two doses of T-DXd well, with no change in body weight or animal appearance.

To validate the results of the neratinib plus T-DXd combination in a second experimental model, we used the human breast cancer PDX, WHIM51, which has the same PIK3CA mutation and a HER2 activating mutation at the neighboring codon, 776, to the V777L mutation present in HP mice (Fig. 4F). Computational modeling of the G776insYVMA mutation showed that it behaves identical to the V777L mutation (10). Similar to the results seen with HP breast cancer organoids, the WHIM51 PDX showed the greatest tumor regression with neratinib plus T-DXd.

The effect of this drug combination was statistically significant as compared to either neratinib or T-DXd monotherapy and compared to the vehicle control (Fig. 4F).

### Proteomics using organoids derived from P and HP mice tumors

In order to characterize the mechanism causing the rapid breast cancer growth in HP mice, we first measured protein phosphorylation using proteomics. Given the differences observed in the HP mice, we examined the key signaling pathways using the Cell Signaling Phospho Antibody Array kit, comparing organoids derived from the P and HP groups (Fig. 5A). The antibody array includes PI3K/AKT signaling, apoptosis, autophagy, cell cycle, ErbB, focal adhesions, MAPK, p53 signaling, VEGF, and other key markers. This array showed a significant increase in the phosphorylation of 18proteins in HP relative to P mice organoids (Fig. 5A-5B). This suggested that H and P mutations cooperate to increase signaling above that achieved with the *PIK3CA* mutation on its own. The increase in cell cycle markers of HP organoids was accompanied by an increased expression of the p-P53, p-P27, and p-PDK1, suggesting that HER2^V777L^ promoted cell proliferation by enhancing the cell cycle process (Fig. 5B). p-AKT, p-GSK3B and p-S6 were also found to be significantly increased (confirmed by Western blot, Supplementary Fig. S1). In summary, the Akt-mTOR-GSK3B signaling axis was significantly altered in HP mice-derived organoids. Further, our data suggests that HER2^V777L^ expression in the mammary gland promoted a continuous enhancement of cell cycle markers, which was consistent with the aggressive rate of growth observed in our transgenic mouse model.

**Fig. 5:**
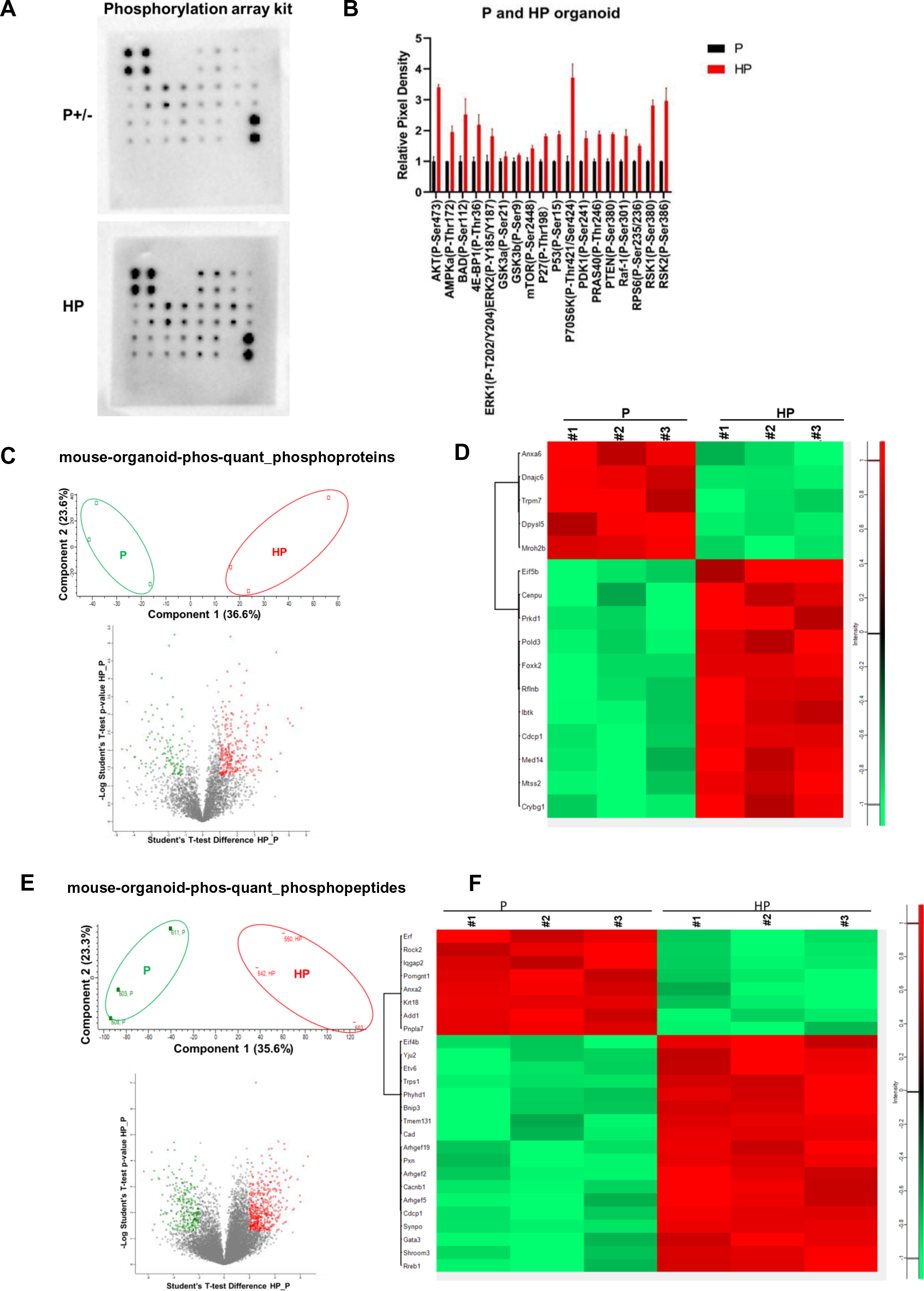
Proteomics using organoid derived from *PIK3CA*^H1047R^ (P), HER2^V777L^; *PIK3CA*^H1047R^ (HP) transgenic mouse tumors. (A) Western blot of Cell Signaling Phospho Antibody Array kit comparing organoids derived from the P and HP groups. (B) Quantitative result of the WB signal intensity of the phospho proteins using the Cell Signaling Phospho Antibody Array kit, comparing organoids derived from the P and HP groups. (C) Top. PCA of the P and HP organoid lysate for quantitative phosphoproteins. Bottom. Volcano plot illustrating phosphoproteins meeting cutoffs for differential expression [log fold change (logFC2) >1, Padj. < 0.05] between P and HP tumors organoids (n = 3 tumors organoid used in the comparison). (D) Heat map showing the clusters of up and down-regulated phosphor proteins (logFC2>1, Padj<0.05) between P and HP organoid samples. Phosphoproteins upregulated in HP tumors organoid are highlighted in red, and phosphoproeins downregulated in HP tumors organoid are highlighted in green. (E)Top. PCA of the P and HP organoid lysate for quantitative phosphopeptides. Bottom. Volcano plot illustrating phosphopeptides meeting cutoffs for differential expression [log fold change (logFC2) >1, Padj. < 0.05] between P and HP tumors organoids (n = 3 tumors organoid used in the comparison). (F) Heat map showing the clusters of up and down-regulated phosphopeptides (logFC2>1, Padj<0.05) between P and HP organoid samples. Phosphopeptides upregulated in HP tumors organoid are highlighted in red, and phosphopeptides upregulated in HP tumors organoid are highlighted in green.

To further elucidate the function of the HER2^V777L^ mutation in the HP mice tumor model, we then performed mass-spectrometry based quantitative proteomics on P and HP breast cancer organoids (Fig. 5C-5H). Organoids were prepared from three independent animals per genotype.

Principle component analysis (PCA) analysis showed these two groups were well-differentiated in both protein and peptide level data (Fig. 5C,5F). Additionally, the divergence analysis result showed the distinct separation of the differentiated protein cluster set (Fig. 5E). Phosphorylation of the cell division markers (e.g., Cenpu, Prkd1), DNA replication and repair markers (e.g., Pold3), and translation machine components (e.g., Eif4b and Eif5b) were increased in HP breast cancer organoids while phosphorylation of the cell senescence marker (e.g., Erf) was significantly decreased (Fig. 5G). Interestingly, Annexin A2, a member of the annexin proteins which belongs to the calcium-dependent phospholipid-binding protein family and plays a vital role in tyrosine kinases / adaptors signal transduction pathways, was increased in P organoids (Fig. 5G). Phosphorylation of Anexin2 is associated with drug resistance in various cancer types (24).

### RNA sequence data using organoids derived from H, P, HP mice tumors and WT mice littermate mammary gland

We next examined gene expression changes using RNA sequencing. We established organoids from H, P, and HP mice tumors and normal mammary glands from WT littermate mice and performed RNA sequencing analysis on 16 independent organoid cell line samples which were established from separate mice. PCA indicated good concordance between the replicates (Fig. 6A). RNA sequencing analysis revealed substantial overlap in differentially expressed genes in single and double mutant samples. 20% of the transcripts that were up-regulated in HP versus WT groups overlapped with both H and P versus WT (Supplementary Fig. S2A). Meanwhile 30% of the significantly downregulated transcripts similarly overlapped between HP versus WT and H or P versus WT (Supplementary Fig. S2B). In comparison, few transcripts were commonly significantly upregulated or downregulated in HP double mutant versus H and P single mutants (Benjamini-Hochberg adjusted p-value < 0.05 and abs(log2FC) > 1.5, Supplementary Fig. S2A, 2B). These data are consistent with the idea that that HER2^V777L^ and *PIK3CA^H1047R^* each affect distinct expression programs that converge when both mutations are present.

**Fig. 6:**
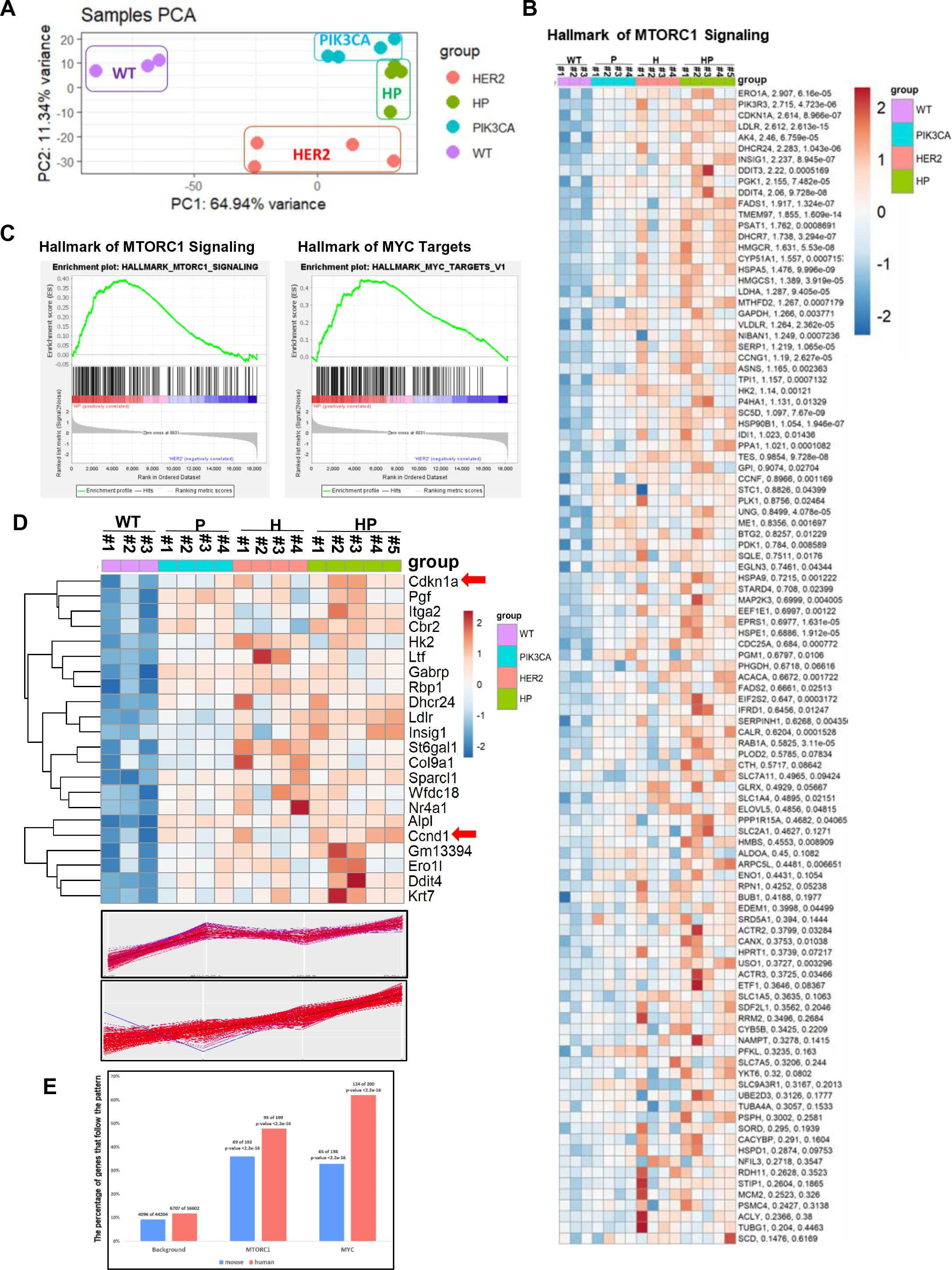
RNA sequence using breast organoid derived from wide type (WT), HER2^V777L^ (H), *PIK3CA*^H1047R^ (P), HER2^V777L^; *PIK3CA*^H1047R^ (HP) transgenic mouse tumors. (A) PCA analysis confirms transgene-based lineage relationship and genetic heterogeneity in RNA samples. (B) Expression of the core enriched genes in the HALLMARK_MTORC1_SIGNALING gene set across all samples. This pathway was identified by GSEA based on the comparing HP and WT. The gene symbol, log2FC of HP versus WT, and Benjamini Hochberg adjusted p-value of HP versus WT are shown to the right. The color key indicates the expression value (normalized and scaled log2 transformed counts): dark blue: lowest; dark red: highest. (C) GSEA results for two hallmark pathways enriched for differentially expressed genes between the HP and H organoid samples. (D) Top, heatmap showing the clusters of DEGs (abs(log2 FC) > 1, Padj < 0.05) between WT, H, P, and HP organoid samples. Bottom, gene expression patterns between WT, P, H and HP groups.

We also observed that there were three times as many differentially expressed genes (DEG) in HP versus H samples (1129 DEGs) relative to HP versus P (340 DEGs). GSEA analysis using the Hallmarks gene set indicated that HP versus H DEGs were significantly enriched for genes related to DNA repair, EMT, and cell cycle related (Supplementary Fig. S2C, 2D). GSEA analysis of differentially expressed genes in HP versus WT revealed many pathways involved in proliferation and cell cycle control further supporting the malignant features of HP model (Supplemental Fig. S2E). These data confirmed the distinct effects of *PIK3CA* mutations and HER2 mutation.

We next examined how genes identified as differentially expressed in one comparison behaved across all the samples. As shown in Supplementary Fig. S3, different pattern clusters of DEGs between WT, H, P, and HP organoid samples emerged including classes of genes that were unique to H samples, and those that were shared between individual mutants (H or P) and double mutants. Of particular interest was the finding that 131 genes followed a stepping pattern such that they were up- (or down-) regulated in both single mutants and then further up- (or down-respectively) regulated in double mutant samples. Consistent with the proteomic data, pathway analysis showed that these genes were enriched for in the PIK3-Akt-mTOR signaling pathway (Fig. 6B). The GSEA analysis showed that the mTOR pathway and the MYC target signature were significantly upregulated in the HP organoid group compared with the H group (Fig. 6C). We also found several critical cell cycle-related markers like *CDKN1A* (which encodes p21) and *CCND1* (Cyclin D1) that are upregulated in single mutant samples and further up-regulated in the HP group (Fig. 6D). Taken together, our data suggested that the co-expression of HER2^V777L^ and *PIK3CA^H1047R^*amplified the signaling transduction cascade from the cell surface receptors to the nucleus causing genetic expression change, resulting in increased oncogene expression and tumorigenesis in our mice.

### HER2 and *PIK3CA* mutations in human patients follow the same signal pathways as in HP mouse model

To understand whether trends observed in mouse tumors are also found in human disease, we next examined the HER2 and PIK3CA mutations in available TCGA RNA-seq data. We examined data available from 113 normal solid tissue from patients, 113 samples with the H1047R PIK3CA mutation, 65 samples with ERRB2 amplification, and 11 samples with both H1047R PIK3CA mutation and ERBB2 amplification. As shown in Supplementary Fig. S4A, the PCA analysis confirms transgene-based lineage relationship and genetic heterogeneity in human mutated samples from -BRCA data. PAM50 classification of these tumors indicated as expected that tumors with HER2 mutations are enriched for Her2 subtypes and tumors with PIK3CA mutations or both HER2 and PIK3CA mutations are enriched for Luminal A and B subtypes (Supplementary Fig. S4B). However, PAM50 subtypes cannot differentiate the samples with HER2 mutations only, *PIK3CA* mutations only, or with bothHER2 and *PIK3CA* mutations (Supplementary Fig. S4B). We next examined whether the expression of genes in the MYC and mTOR pathways followed a similar stepping pattern between individual and double mutations as found in mouse samples. In TCGA samples, 95 out of 205 genes (46%) in the mTOR and 116 out of 188 genes (62%) in the Myc pathway followed the stepping pattern (Fig.6E. This is significantly enriched (p < 10^-15^, exact binomial test) based on the background distribution of 12% of genes found with the stepping pattern. These data indicated that the finding from our mouse models are consistent with what is observed in human patients.

### p21 mediated cell cycle stimulation in HP mice-derived organoid

To further understand the mechanism of our mouse tumor model, we confirmed our RNA-seq analysis data (Fig. 6D) using western blotting detection on WT, P, H, and HP organoids. The protein level of p21 (also known as p21WAF1/Cip1, and encoded by *CDKN1A*) is upregulated in HP organoid groups (Fig. 7A). p21 can impact apoptosis as well as cell cycle progression in response to many stimuli (25). We found that apoptosis was decreased in P and HP organoids (Fig. 7B, Supplementary Fig. S5A). While p21 can function as an inhibitor of cyclin-dependent kinases, p21 also stabilizes the CCND1-CDK4/6 complex to further activate CDK4/6 (26), allowing the cell to exit the G1 phase and initiate DNA replication. To measure cell cycle changes, we conducted BrdU and Propidium iodide (PI) staining assay on the organoids. As shown in Fig. 7B and Supplemental Fig. S5B, the percentage of BrdU positive cells increased in HP mice derived organoids compared with WT, P, and H mice derived organoids. We next used the CDK4/6 inhibitors abemaciclib and palbociclib to inhibit cell cycle arrest and to further inhibit G1/S checkpoint exit in HP mice-derived organoids. As shown in Fig. 7C, abemaciclib treatment decreased the number of BrdU positive cells. Since abemaciclib has overlapping GI toxicities with neratinib, we tested the combination of neratinib and palbociclib. As shown in Fig. 7D, the *in vitro* treatment of neratinib and palbociclib combination reduced the number of BrdU positive cells relative to palbociclib or neratinib monotherapy. To test this combination *in vivo*, we treated the luciferase labeled HP organoid transplanted mice with palbociclib and neratinib. As shown in Fig. 7E-G, the HP organoid transplanted tumors grow more slowly in neratinib plus palbociclib treatment group than in either monotherapy group. Further, the palbociclib monotherapy group showed drug resistance after 2 weeks treatment, with rapid tumor growth after day 28 (Fig. 7E). Based on both luciferase signal and tumor mass, the neratinib plus palbociclib treatment group showed statistically significant differences compared to either monotherapy arm and to the vehicle control (Fig. 7F-G). To ensure that the mouse received an adequate dose of palbociclib, WBC and neutrophil counts were measured. The palbociclib treated mice, both in the palbociclib monotherapy and combination therapy groups, had a decrease in neutrophil count (Supplementary Fig. S6C), matching the effect of palbociclib on patients. To validate the results of the neratinib plus palbociclib combination in a second experimental model, we again used the WHIM51 PDX model (Fig. 7H-I). Similar to the results seen with HP breast cancer organoids, the WHIM51 PDX showed the greatest tumor regression with neratinib plus palbociclib and this effect was statistically significant as compared to either neratinib or palbociclib monotherapy and compared to the vehicle control (Fig. 7I). In conclusion, these data suggest combining HER2 and CDK4/6 inhibition as a treatment strategy for tumors containing HER2 and PIK3CA mutations.

**Fig. 7:**
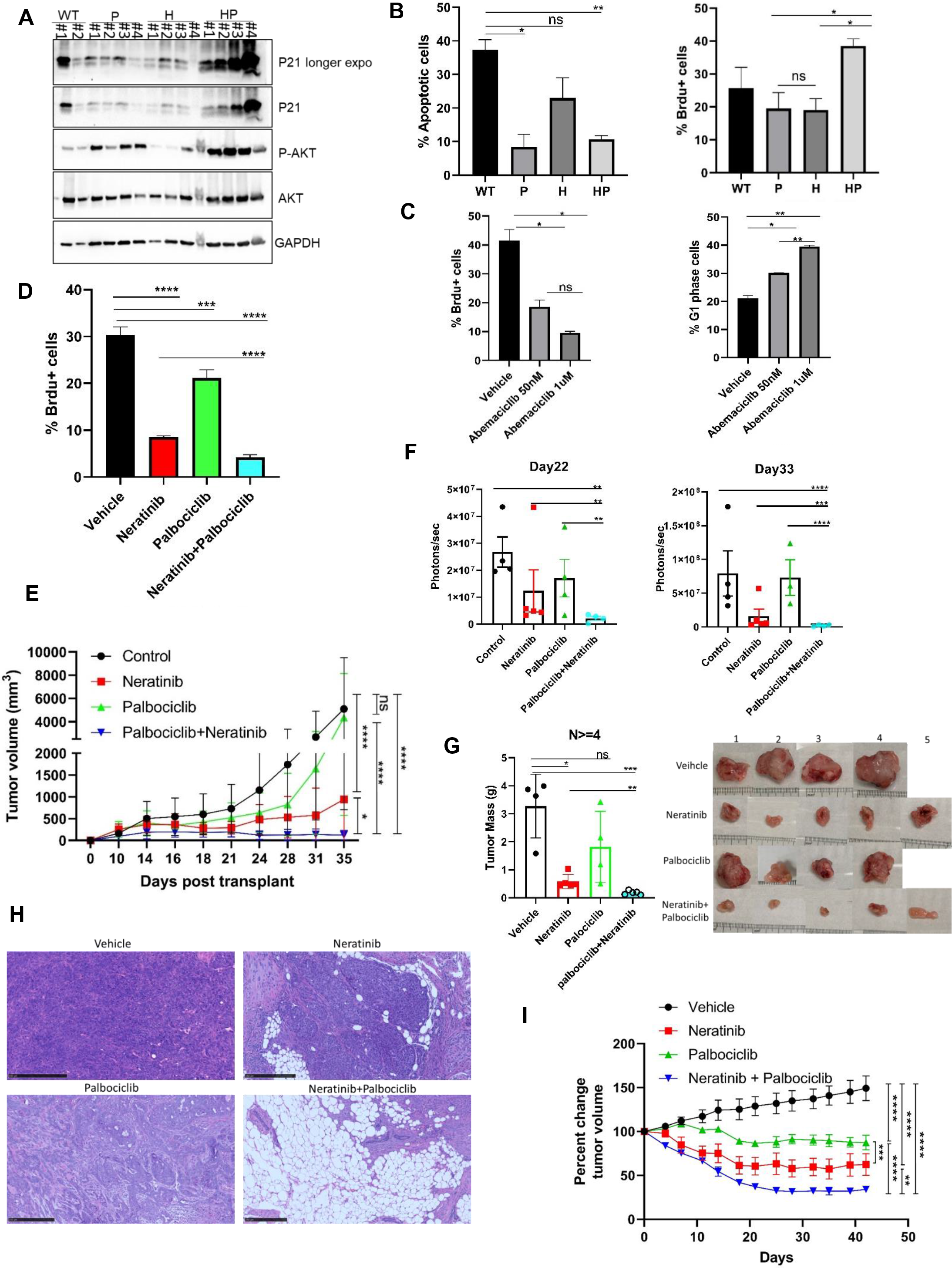
P21 mediated cell cycle stimulation in HP mice-derived organoids. (A) Western blotting P21, phospho-AKT and total AKT, and GAPDH from indicated mice breast derived organoids. (B) Flow cytometry assay on organoids from WT, H, P, HP mice breast using AnneinV and Propidium iodide (PI). Quantification of the percentage of apoptotic organoid cells from WT, H, P, HP mice group. (C) Flow cytometry assay on organoid from WT, H, P, HP mice breast using BrdU and Propidium iodide (PI) staining kit. Quantification of the percentage of BrdU positive organoid cells from WT, H, P, HP mice group. (D) Flow cytometry assay on HP organoid treated with neratinib, palbociclib and combination of neratinib and palbociclib using BrdU and Propidium iodide (PI) staining kit. Quantification of the percentage of BrdU positive organoid cells from untreated and treated groups. (E) HP breast tumor organoid cells (1 × 10^6^) were orthotopically injected into the 8-10 week-old female FVB x C57BL/6J F1 (FVBB6F1) mice (n = 5–6). On day 4, starting the treatment with saline water, neratinib only, palbociclib only, and neratinib combine palbociclib respectively for 32 days. Tumor volume growth rate of treated mice in 4 arms are shown in the Figure. (F) BLI intensity measurements on day22 and day33. (G) Left, Transplant mice tumors were collected on day36 and tumors weight was measured. Right, tumor images of treated mice upon collecting. (H) Representive hematoxylin and eosin (H&E) staining image of the tumor slides from transplant mice within treatment. Scale bars are 250 µm. (I)WHIM51 tumor cells (3.3 × 10^6^) were injected into the 8–10-week-old female NSG mice (n = 5–6). When tumor size reached 300mm^3^, treatment with saline water, neratinib only, palbociclib only, and neratinib combine palbociclib was started. Tumor volume change percentage of treated mice in 4 arms as shown in the Figure. Data are plotted as means± SEM, 1 dot represents 1 data point. ∗P < .05, ∗∗P < .01, ∗∗∗P < .001, and ∗∗∗∗P < .0001 as calculated by the Mann-Whitney U test and two-way ANOVA group test.

## Discussion

HER2 activating mutations were identified in breast cancers that lack HER2 gene amplification (3), and we have previously shown that patients with HER2 mutated, metastatic breast cancers respond to the HER2 tyrosine kinase inhibitor, neratinib (7). While combining neratinib with other breast cancer drugs, such as fulvestrant or trastuzumab, improves response rates and duration of response, there is concern that mutations in downstream signaling proteins, such as PIK3CA, can reduce response to neratinib. We found that HP transgenic mice develop aggressive breast cancers that metastasize to the lungs and are resistant to neratinib. Testing other targeted therapies showed that T-DXd was a highly effective drug for HP breast cancers, whereas trastuzumab on its own had no effect and the PIK3CA-specific inhibitor, alpelisib, had only a modest effect. Combining neratinib plus T-DXd showed statistically significant reductions in tumor size in both HP mice breast cancers and in a human breast cancer PDX, WHIM51, than either drug on its own. Proteomic and RNA-Seq analysis of HP breast cancers showed increased gene expression of p21WAF1/Cip1 and Cyclin D1 (CCND1), which prompted us to combine neratinib with CDK4/6 inhibitors. We found that neratinib plus palbociclib gave a statistically significant reduction in mouse HP breast cancers and human WHIM51 PDX as compared to either drug alone.

p21WAF1/CIP1 plays a dual role in cancers, with both tumor suppressor and oncogenic functions(27). While p21 can cause cell cycle arrest in response to DNA damage(28), it also stabilizes the CDK4/6-CyclinD1 complex to promote the G1-S phase transition (29,30). Mouse embryo fibroblasts lacking both p21 and the related gene, p27/kip1, fail to assemble detectable amounts of the cyclinD1-Cdk complex and are unable to direct CyclinD1 into the nucleus(29). Sequestration of p21 by CDK4/6-CyclinD1 may promote cellular proliferation by freeing CDK2 and PCNA from the inhibitory effects of p21 (31). The cell-cycle entry program starts with activating CDK4/6. CDK4/6 activity increases rapidly before CDK2 activity gradually increases (32). The abundance of p21 maintains the ratio of the p21-cyclinD1-CDK4/6 axis to further improve cell survival and cell proliferation. This leaves open a question of how p21 abundance is properly regulated in HP tumor cells. The binding of p21 to cyclinD1 prevents its export from the nucleus and degradation in the cytoplasm (33). This suggests that cyclinD1 and p21 stabilize each other in HP tumor cells. There is a controversy as to whether CDK4/6 inhibitors like palbociclib cause cell cycle arrest by p21-dependent or independent mechanisms (34,35), therefore, we targeted the p21-cyclinD1-CDK4/6 axis instead of p21 itself to avoid this controversy. In our study, we show the p21-cyclinD1-CDK4/6 axis promotes cell cycle entry in HP tumor cells, demonstrating the oncogenic role of p21 in the HP breast cancer model.

Hyperactivation of the PI3K-AKT-mTOR pathway mediates drug resistance and tumor progression (36). Activation of the PI3-kinase pathway either by loss of PTEN or activating mutations in PIK3CA cause resistance to trastuzumab (37,38). PI3-kinase hyperactivation results in lapatinib resistance (39). PIK3CA mutations are found in both HER2-amplified breast cancer (12) and HER2-mutated breast cancer (5,7). A lower pathological complete response rate to neoadjuvant trastuzumab plus lapatinib in HER2-amplified breast cancers that had PIK3CA activating mutations was observed (12). Currently, clinical trials of drugs targeting the PI3K-AKT-mTOR pathway combine them with endocrine, CDK4/6i, or HER2i therapy to overcome drug resistance, such as ClinicalTrials.gov Identifiers: NCT04862663 and NCT04862663.

While PIK3CA mutations cause resistance to trastuzumab and lapatinib, there is evidence that these mutations may be sensitive to HER2 ADCs. PIK3CA mutations did not affect clinical outcomes with T-DM1 treatment in the EMILIA phase 3 clinical trial (40). Further, in cell line and PDX models, PIK3CA mutations did not cause resistance to T-DM1 (40). The effect of PIK3CA mutations on T-DXd treatment is unknown. As shown in Fig. 4, T-DXd is effective either alone or in combination with neratinib in HP tumor cells, suggesting T-DXd will benefit PIK3CA mutant, HER2-activated patients. Clinical trials of pan-PI3K inhibitors with HER2-targeted therapies have been unsuccessful because of toxicity (41). While newer trials are looking at the PIK3CA-specific inhibitor alpelisib or second-generation mTOR inhibitors like sapanisertib, toxicity concerns remain and therefore we explored other drugs that can be combined with neratinib (42). In our study, we combine neratinib with both T-DXd and palbociclib to overcome drug resistance, and our data have shown effective pre-clinical results. Two phase I, clinical trials are currently testing the safety and efficacy of neratinib with T-DXd (ClinicalTrials.gov Identifier: NCT05372614 and NCT05274048) and another phase I clinical trials is combining neratinib and palbociclib (ClinicalTrials.gov Identifier: NCT03065387). Our study provides further pre-clinical evidence supporting these phase I trials and the results of these trials is eagerly awaited.

## Materials and Methods

### Mice

HER2^V777L^ transgenic mice were generated using TALEN-based genome editing as previously reported (4). *PIK3CA^H1047R^* mice were purchased from Jackson Laboratory, strain #:016977. All animal studies were approved by the Institutional Animal Care and Use Committee (IACUC) at Washington University in St. Louis.

### Induction of Cre-mediated recombination

To study the contribution of double mutation of HER2 and *PIK3CA*, HER2^V777L^ mice were crossed with *PIK3CA^H1047R^* mice to generate HP mice. All experiments were done using syngenic mice (F1 C57BL/6J: FVB mice) and all experiments were performed with littermate control. For adenovirus Cre mediated recombination, 8-10 weeks old mice were injected orthotopically into the mammary fat pad with adenovirus Ad5CMVCre, which was purchased from the Viral Vector Core Facility of the University of Iowa, College of Medicine (titer range 1x10^10 to 7x10^10 pfu/ml).

### *Ex vivo* single drug sensitivity assay

Organoids were removed from Matrigel using Cell Recovery Solution (Corning), washed, and resuspended in media with 6% Matrigel (GFR). 25ul of organoids in 6% Matrigel are added to a 384-well black wall, clear bottom, and tissue culture plate and incubated overnight at 37 degrees C, 5% CO2. DMSO-based drugs are serially diluted 3-fold in 100% DMSO and then diluted into mouse breast tumor media in a 96 well plate at 2X the final desired concentration. Aqueous drug serial dilutions are made directly in media in the 96-deep well plate. The drugs, diluted in mice tumor media, are tested in duplicate with 25ul of drug added to the cells. Control wells are also prepared and added with a vehicle in the media to serve as 100% viability controls. Wells without cells serve as 0% viability controls. Organoids and drugs are incubated for six days. Viability is measured by adding 25ul of Cell Titer Glo 3D (Promega) to each well. Sealed plates are shaken for 5 minutes and incubated for another 30 to 60 minutes at room temperature (signal stable for ∼4 hours). Luminescence was measured in a Tecan plate reader.

### *Ex vivo* drug combination assay

Organoids are removed from Matrigel with Cell Recovery Solution and digested with TrypLE. After washing with Ad DME/F12 media, cells are diluted into cold mouse breast tumor media plus 6% Matrigel (GFR) and 20ul is added per well of a 384 well black wall, clear bottom, tissue culture treated plate. The plate is incubated at 37 degrees C and 5% CO2 overnight. DMSO-based drugs are first serially diluted in 100% DMSO and then diluted into mouse breast tumor media in a 96-deep well plate. Aqueous drug serial dilutions are made directly in media in the 96-deep well plate. The two drugs to be compared are prepared at 3X the final desired concentration and the serial dilutions are 2-fold. Using the adjustable spacing Integra Voyager pipettor, 20ul of each of the two drugs are transferred from the 96 well intermediate drug dilution plate and added to the 384 well assay plate in a grid format as described in Griner, et al. Each drug is also assayed as a single drug with 20ul media making up for the lack of a 2nd drug. Additional wells are seeded with cells as vehicle controls for the 100% viability and wells without any cells for the 0% viability controls. Organoids and drugs are incubated for six days. Viability is measured by adding 30ul of Cell Titer Glo 3D (Promega) to each well and incubating as described. The percent viability for both assays is determined by first calculating the average vehicle value and the average “no cell” value. Then, percent viability is determined with the formula: Percent viability equals 100 minuses (ave viability control minus value)/(ave viability control minus ave no-cell control) X 100. IC50 values and graphs are calculated using GraphPad Prism 5 for Windows. The Synergy mapped to D-R (Loewe) plots are prepared using the freeware program Combenefit 2.021 for Windows from the University of Cambridge, Cancer Research UK.

### Transwell migration and invasion assay

In the transwell cell migration assay, 24-well inserts from Corning were used. Organoid cells were digest into single cell using trpLE and seeded in top of the transwell membrane in the upper chamber in the 100ul DMEM/F12 medium without serum. 600μl of DMEM/F12 medium with 10% FBS was added to the lower chamber. For invasion assay, 10% matrigel was added to the top membrane. Cells in the bottom was fixed with 4%PFA after 3days and stained with 0.02% crystal violet. The number of migrated or invaded cells could be quantified by counting underneath a microscope or in the pictures taken.

### Wound healing assay

In the wound healing assay, 96-well plate from Corning were used. Organoid cells were digest into single cell using trpLE and seeded in the plate. Wound were generated by AccuWound 96 scratching tool, images were taken every 3 hours by incucyte S3 for 48 hours.

### *In vivo* therapy

Drug treatments are conducted on three different HP cancer models. 10-12weeks old female HP mice with orthotopic Adenovirus-Cre injection on the mammary gland or PDX WHIM51 are randomized to 4 treatment groups (n=3∼6 per group): 1) vehicle, 2) neratinib, 3) T-DXdonly (2 dosages for 2 weeks), and the 4) neratinib plus T-DXd. Mice are subjected to the treatment from the 4th-day post-injection. This drug combination is chosen because we have already determined that HP mouse breast tumors are neratinib resistant, but sensitive to T-DXd.

Female P or HP mice at 8-12 weeks old are randomized to 4 treatment groups (n=3∼6 per group): 1) vehicle, 2) alpelisib, 3) T-DXd only (2 dosages for 2 weeks), and 4) alpelisib plus DS8201. HP transplant mice or PDX WHIM51 subcutaneously implanted mice are randomized to 4 treatment groups (n=3∼6 per group): 1) vehicle, 2) neratinib, 3) palbociclib, and 4) neratinib plus palbociclib. Neratinibis dosed at 40 mg/kg daily by oral gavage to animals. T-DXd is dosed at 30 mg/kg weekly by tail vein injection to animals. Palbociclib is dosed at 62.5 mg/kg weekly by oral gavage to animals. Alpelisib dosed at 25mg/kg daily by oral gavage to animals. All animal experiments are conducted in compliance with federal guidelines and approved by the institutional animal care and use committee.

### *In vitro* therapy

*In vitro* experiments: HP breast cancer organoids and PDX WHIM51 human breast cancer organoids are routinely cultured in 3D culture with matrigel as previously described (43,44)). Drug concentration added as indicated.

### Immunohistochemistry and Immunofluorescence of paraffin-embedded sections

Tumor tissues are fixed with 10% formalin, embedded in paraffin, and cut into 5-µm sections (Digestive Diseases Research Core, Washington University School of Medicine). Sections are de-paraffinized, hydrated, and treated with heat-activated antigen unmasking solution (Vector Laboratories). Immunostaining is performed using the antibodies listed in the table below. For immunofluorescence staining, antibody binding is visualized with Alexa Fluor 488 or 555 or 647 fluorochromes, then counterstained with DAPI-containing mounting media (Sigma). For immunohistochemistry staining, DAB substrate (Cell Signaling) is applied, then counterstained with hematoxylin (Thermo Fisher Scientific). Primary antibodies used in the study include rabbit ERBB2 (CST, 2165S), mouse ERBB2 (CST, AF2967-SP), Smooth Muscle Actin Antibody (Santa Cruz Biotechnology, SC-53142), Anti-FOXA1 Antibody (Abcam, ab173287), Era Antibody (D-12) (Santa Cruz Biotechnology, SC-8005), GATA-3 Antibody (Santa Cruz Biotechnology, SC-268), Cytokeratin 14 Antibody (Santa Cruz Biotechnology, SC-53253), Cytokeratin 8/18 Antibody (Fitzgerald Industry International, 20R-CP004), phospho-p44/42 MAPK (Erk1/2) (Thr202/Tyr204) (Cell signaling technology, #9101). Secondary antibodies used include anti-rabbit IgG (H+L) Alexa Fluor 555 (Invitrogen, A-31572), anti-goat IgG (H+L) Alexa Fluor Plus 488 (Invitrogen, A32814), Goat anti guinea pig (H + L) FITC (Fitzgerald Industry International, 43R-1095), SignalStain Boost IHC Detection Reagent (HRP, Rabbit) (Cell signaling technology, 8114S).

### Establishment of breast organoid

Isolation of epithelial organoids from the mammary gland of wide type, or tumor from HER2^V777L^ only, *PIK3CA^H1047R^* only, HER2V777L; *PIK3CA^H1047R^* transgene mice as previously described (43). Isolated crypts were embedded in Matrigel (BD Biosciences) and seeded in 6-well plates.

The cells were overlaid with 2 mL/well basal culture medium (advanced Dulbecco’s modified Eagle medium/F12 supplemented with penicillin/streptomycin, 10 mmol/L HEPES, Glutamax, 1 × B27 [all from Thermo Fisher Scientific], 125 µM N-acetyl-cysteine (Sigma), 50 ng/ml murine epidermal growth factor (EGF, Invitrogen), 10% Rspo1-Noggin-conditioned medium (Noggin and R-spondin combined expression 293T cell line to generate conditioned media were a gift from Blair Madison). Organoids from healthy mammary tissue were isolated in a similar manner and cultured as described previously (44), with modifications. Healthy organoids were cultured in AdDMEM/F12 supplemented with 1 M HEPES, Glutamax, penicillin/streptomycin, 1% insulin-transferrin-selenium (vol/vol, Gibco) and 2.5 nM FGF2 (Sigma).

### Organoid transplant mice model

Organoid cells were cultured as previously described. We collected organoid cells and digest the cells with TripLE for 5-10 minutes. The cells were counted and 1x10^6^ Organoid cells were injected in the fourth mammary fat pad of wild type of syngenic mice (F1 C57BL/6J: FVB mice). Tumors were treated with a drug on the fourth day and collected after 26 days post-injection.

### *In vivo* BLI

For BLI of live animals, mice were injected intraperitoneally with 150 μg /g D-luciferin (Gold Biotechnology, St. Louis, MO) in PBS, anesthetized with 2.5% isoflurane, and imaged with a charge-coupled device (CCD) camera-based system (IVIS 50, PerkinElmer, Waltham, MA; Living Image 4.3.1, exposure time 10-60 seconds, binning 8, field of view 12cm, f/stop 1, open filter, ventral view). Luminescence was displayed as photons/sed/cm2/sr. Regions of interest (ROI) were defined manually over the lower abdomen using Living Image 2.6 with measurements reported as photons/sec.

#### Pearl Image

NIR-trastuzumab was injected into the tail vein of H or HP transplant mice bearing breast tumor one day prior to imaging. The near-infrared imaging was taken using the Pearl Trilogy (LI-COR, Lincoln, NE) in the Washington University Molecular Imaging Center (MIC). The PearlCam and ImageStation software was used for the measurement and analysis.

#### RNA-seq

Base-calling and demultiplexing were performed with Illumina’s bcl2fastq software (v2.20) with a maximum of one mismatch in the indexing read. RNA-seq reads were then aligned to the Ensembl release 96 top-level assemblies with STAR (v2.0.4b)(45). Gene counts were derived from the number of uniquely aligned unambiguous reads by Subread:featureCounts (v1.4.5)(46). Sequencing performance was assessed for the total number of aligned reads, the total number of uniquely aligned reads, and features detected. The ribosomal fraction, known junction saturation, and read distribution over known gene models were quantified with RSeQC (v2.3)(47). All the gene counts were imported into the R/Bioconductor package DESeq2 (v1.33.5)(48). Mutation groups were modeled as the variates for differential gene expression testing. The raw counts were normalized with the median of ratios method provided by DESeq2 and effect size shrinkage was performed using the ashr method.

#### Gene set enrichment analysis

Whole DESeq2-normalized counts were subjected to gene set enrichment analysis using GSEA (v4.2.2)(49). GSEA comparisons between different mutation groups were performed using previously defined Hallmark gene sets and curated gene sets. Mouse_ENSEMBL_Gene_ID_Human_Orthologs_MSigDB.v7.5.1.chip was used as annotation file. The permutation type was gene_set and the number of permutations was 1000.

#### Gene expression pattern clustering

K-means clustering was performed using the Z-score on DESeq2-normalized counts data of significant genes. Differentially expressed genes (Benjamini-Hochberg adjusted p-value < 0.05 and abs(log2FC) > 2) from any comparison (WT vs H, WT v P, WT vs HP, H vs HP, P vs HP, and H vs P, 1572 total genes) were divided into 25 clusters. All results were calculated and visualized in R using stats (v4.1.0), ggplot2 (v3.3.5), and pheatmap (v1.0.12).

#### TCGA RNA-seq analysis

RNA-Seq data from the TCGA-BRCA project were downloaded from Genomic Data Commons Data Portal (v31.0) using the package TCGAbiolinks (v2.22.1)(50). H1047R mutated samples were chosen through the GDC Exploration page and ERBB2 amplified samples were those annotated as amplified based on the CNA field in cBioPortal. Using these criteria, we identified 113 normal samples, 106 single H1047R mutated samples, 82 single ERBB2 amplificated samples, and 12 double mutated samples. PAM50 subtype assignments are from the cBioPortal. Human genes annotation was performed using the org.Hs.eg.db (v3.14.0) Bioconductor package. The Ensembl gene ids between humans and mice were transformed using biomaRt (v2.50.3).

#### Proteomics Analysis

All sample data were analyzed using the Perseus software (version 1.6.15.0). The intensities of proteins were extracted from Proteome Discoverer v2.3 search results and log-based 2 transformed. The abundances were categorized into 2 different categories: control and disease. The imputation for the missing abundances was performed by assigning small random values from the normal distribution with a downshift of 1.8 SDs and a width of 0.3 SDs. All abundances for each sample were further normalized by subtracting the median from each sample abundances. Then, the unpaired two-tail student t-test was performed, and the difference in averages was calculated for the comparison between disease and control. Volcano plots were created for the comparison with cut-off values of FDR = 0.05 (-log10(0.05) = 1.30) and log base 2 difference = 2, which equals to a 4-fold-change. The PCA plot was created for all phosphopeptides/phosphoproteins. The heatmap was created for all phosphopeptides/phosphoproteins with FDR less than 0.05.

## Statistical analysis

Data are presented as means and SEM. The Student *t-*test and the Mann-Whitney U test were used to analyze data in 2 groups. For multiple groups, 2-way ANOVA analysis of variance with appropriate posttests were used. For statistical analysis, GraphPad Prism (version 8; GraphPad Software, Inc, La Jolla, CA) was used. A *P* value less than .05 was considered statistically significant. All authors had access to the study data and reviewed and approved the final manuscript.

## Abbreviations

EGFR: Epidermal Growth Factor Receptor
HER2: Human Epidermal Growth Factor Receptor 2
PIK3CA: Phosphatidylinositol-4,5-Bisphosphate 3-Kinase Catalytic Subunit Alpha
TCGA: The Cancer Genome Atlas
ADCs: Antibody-drug conjugates
T-DXd: Trastuzumab deruxtecan
CDK4/6i: Cyclin-dependent kinase 4 and 6 inhibitors.

## Data availability

RNA sequencing data generated in this study have been deposited in the Gene Expression Omnibus with accession codes GSE216871. The dbGaP Study Accession number for TCGA data is phs000178. All other data supporting the findings of this study are available from the corresponding author upon reasonable request. Source data are provided in this paper. All the analyses were based on standard algorithms described in the Methods and referenced accordingly. There are no custom algorithms to make available.

---

**Supplemental Fig. S1:**
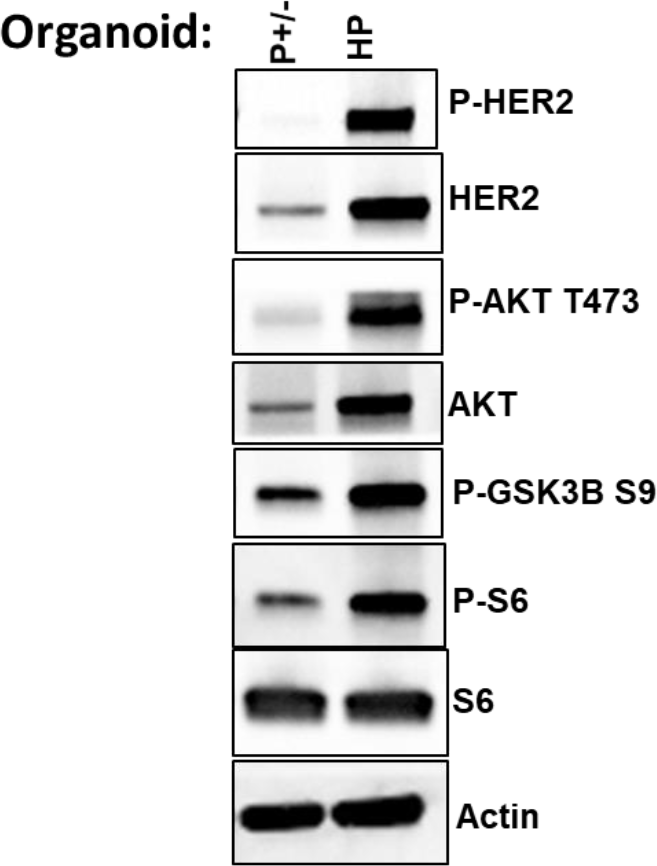
Western blot of Phospho AKT, S6, GSK3B and total AKT, S6, Actin signal comparing organoids derived from the P and HP groups.

**Supplemental Fig. S2:**
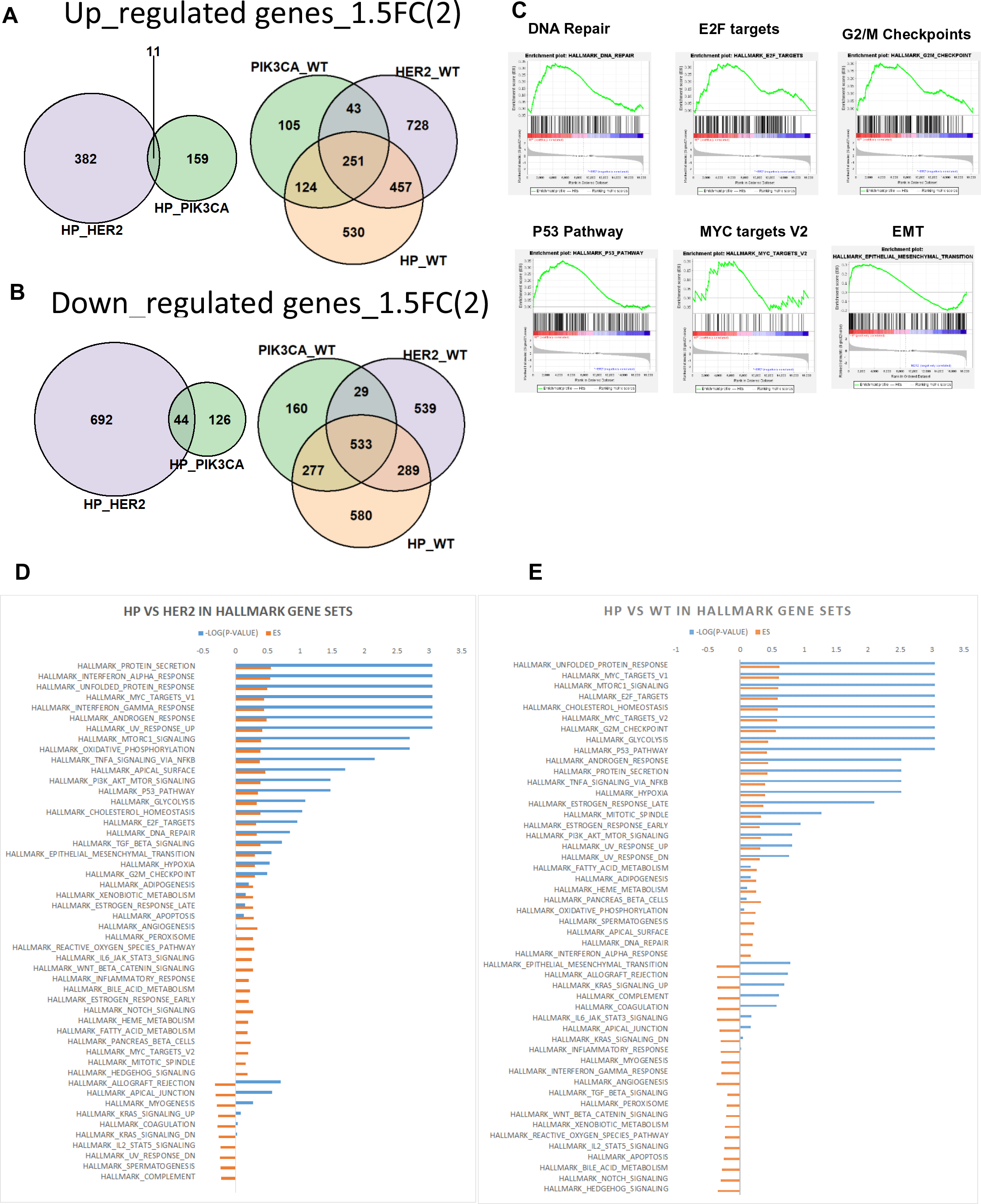
(A) Venn diagrams showing the number of genes up-regulated in the comparisons between HP versus HER2 and HP versus PIK3CA (left), and the comparisons between PIK3CA versus WT, HER2 versus WT, and HP versus WT (right). (B) Venn diagrams showing the number of genes down-regulated in the comparisons between HP versus HER2 and HP versus PIK3CA (left), and the comparisons between PIK3CA versus WT, HER2 versus WT, and HP versus WT (right). (C) Representative Hallmark Gene sets significantly enriched in HP compared with H as identified by Gene set enrichment analysis (GSEA, p < 0.05)). ES (enrichment score) and -log10(p-values) of pathways are shown. (D) Gene sets significantly enriched in HP compared with H as identified by GSEA (p < 0.05)). ES (enrichment score) and -log10(p-values) of pathways are shown. GSEA was performed using the Hallmark gene sets in the Molecular Signatures Database (v7.5.1). (E) Gene sets significantly enriched in HP compared with WT as identified by GSEA (p < 0.05). ES (enrichment score) and -log10(p-values) of pathways are shown from GSEA performed using Hallmark Gene sets in the Molecular Signatures Database (v7.5.1).

**Supplemental Fig. S3:**
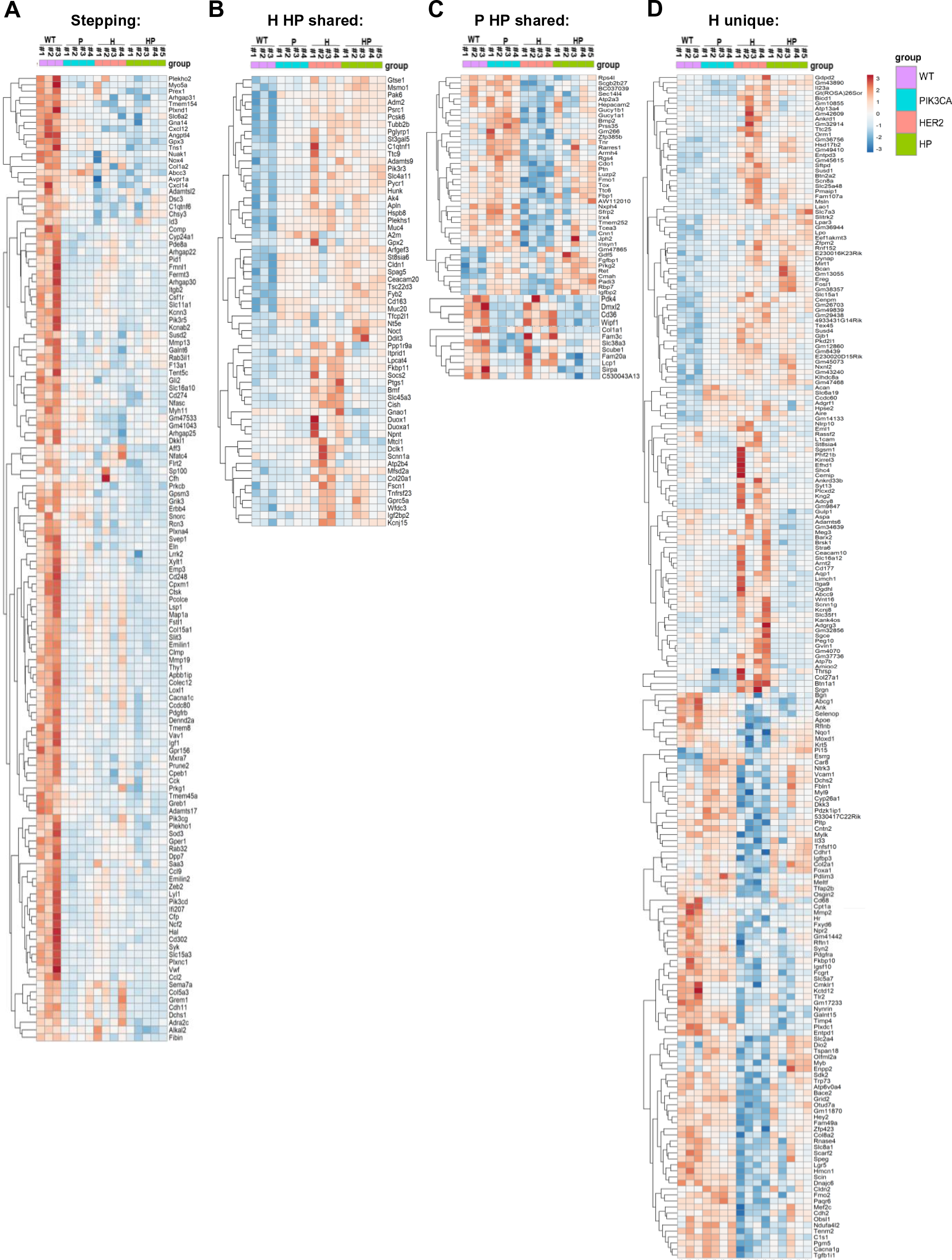
Expression signature analysis. (A) Heat map showing the stepping pattern clusters of DEGs (log2 FC > 1, Padj < 0.05) between WT, H, P, and HP organoid samples. (B) Heat map showing the H and HP shared pattern clusters of DEGs (log2 FC > 1, Padj < 0.05) between WT, H, P, and HP organoid samples. (C) Heat map showing the P and HP shared pattern clusters of DEGs (log2 FC > 1, Padj < 0.05) between WT, H, P, and HP organoid samples. (D) Heat map showing the H unique pattern clusters of DEGs (log2 FC > 1, Padj < 0.05) between WT, H, P, and HP organoid samples.

**Supplemental Fig. S4:**
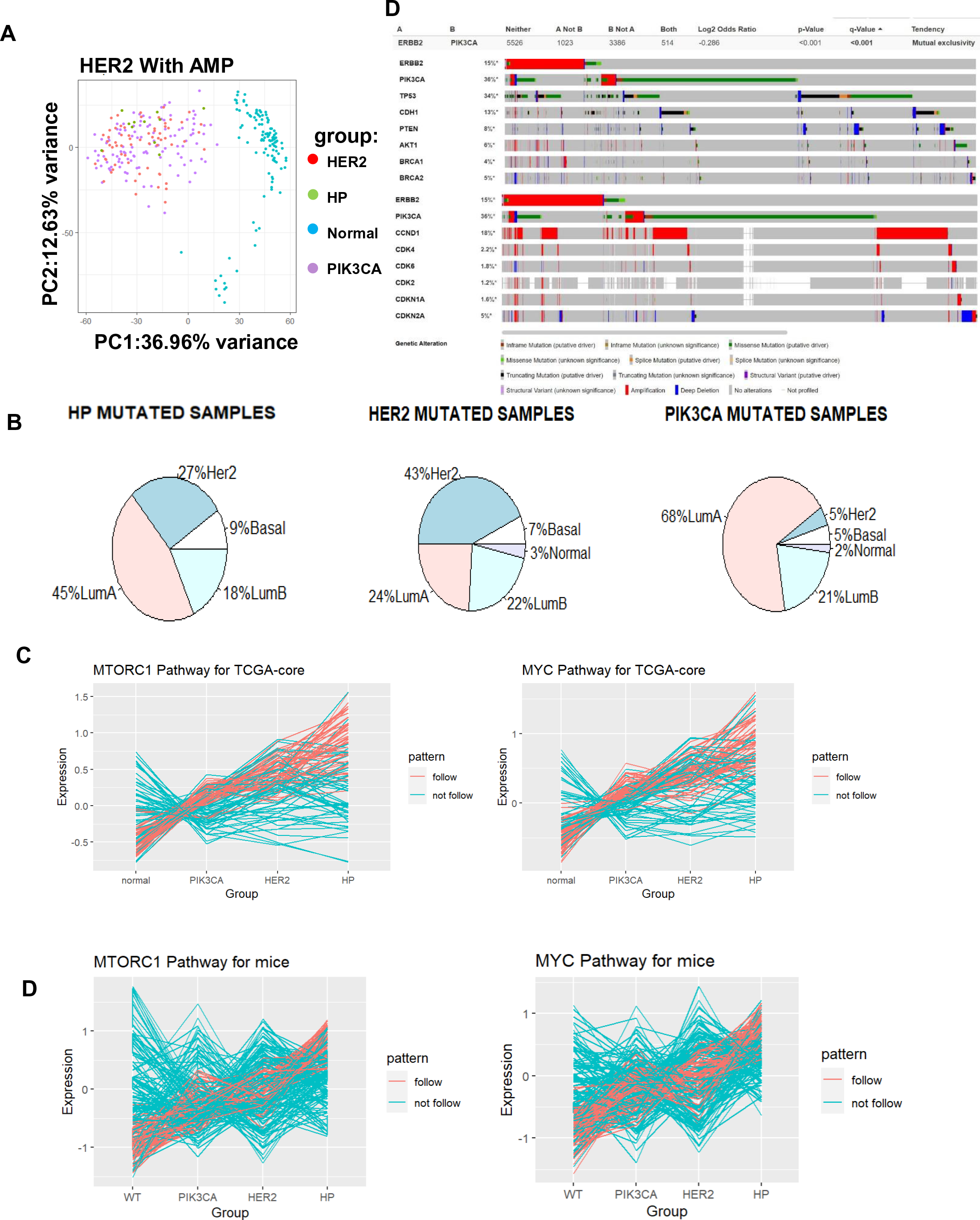
(A) Left, PCA analysis confirms transgene-based lineage relationship and genetic heterogeneity in human mutated samples from -BRCA data (HP mutated: n=11, HER2 mutated: n=65, PIK3CA mutated: n=113). Right, Significant P value of mutual exclusively between HER2 and PIK3CA from TCGA. (B) Relationship of PIK3CA mutation and HER2 mutation and PAM50 subtypes based on annotations from the TCGA-BRCA data (HP mutated: n=11, HER2 mutated: n=65, PIK3CA mutated: n=113). (C-D) Samples in TCGA data with PIK3CA, Her2, and both mutations follow a similar step pattern in MYC and MTORC1 pathways as observed in both human and mice. Line charts showing the average gene expression pattern for each gene in MYC pathway and MTORC1 pathway based on TCGA-BRCA mutation annotations. Red lines indicate the expression pattern follows the step expression pattern found in mice. Blue lines indicate other expression patterns. Both MTORC1 and MYC pathways are enriched for genes that follow the step pattern (p < 10^-15^, exact binominal test) relative to the background distribution of expression patterns.

**Supplemental Fig. S5:**
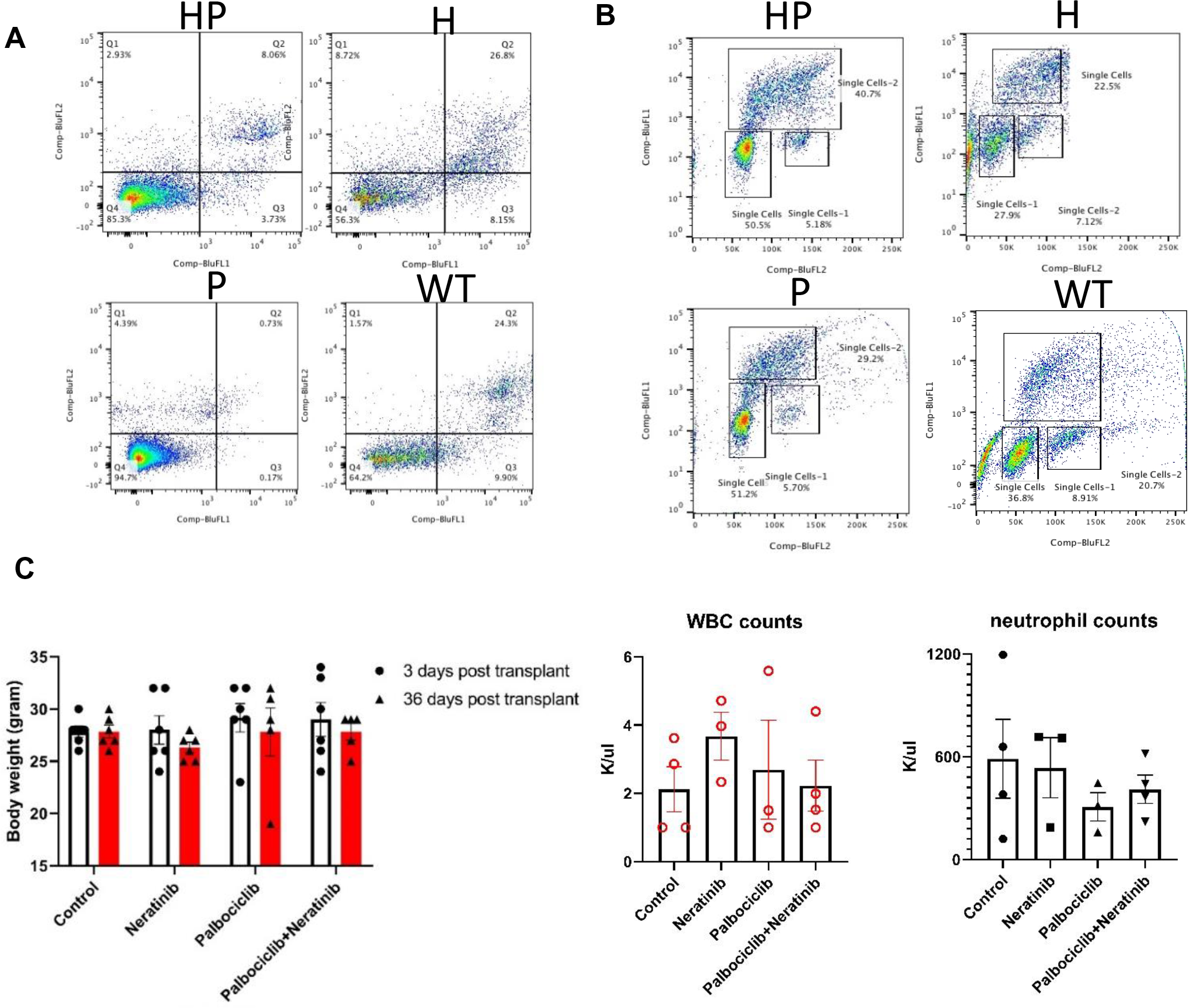
(A) Flow cytometry analysis image on organoids from WT, H, P, HP mice breast using AnneinV and Propidium iodide (PI). (B) Flow cytometry analysis image on organoids from WT, H, P, HP mice breast using BrdU and Propidium iodide (PI) staining kit. (C) Left, Body weight measurements on HP organoid transplant mice in 4 arms treatment groups. Right, Quantification of whole blood cell and neutrophil cell counts after treatment on day36. Data are plotted as means± SEM, 1 dot represents 1 data point.

